# Evaluation of Na_v_1.8 as a therapeutic target for Pitt Hopkins Syndrome

**DOI:** 10.1101/2022.04.15.488505

**Authors:** Keri Martinowich, Debamitra Das, Srinidhi Rao Sripathy, Brady J. Maher

**Author notes:** Correspondence to: Brady J. Maher.

## Abstract

Pitt Hopkins Syndrome (PTHS) is a rare syndromic form of autism spectrum disorder (ASD) caused by autosomal dominant mutations in the Transcription Factor 4 (*TCF4*) gene. *TCF4* is a basic helix-loop-helix transcription factor that is critical for neurodevelopment and brain function through its binding to cis-regulatory elements of target genes. One potential therapeutic strategy for PTHS is to identify dysregulated target genes and normalize their dysfunction. Here, we propose that *SCN10A* is an important target gene of *TCF4* that is an applicable therapeutic approach for PTHS. *Scn10a* encodes the voltage-gated sodium channel Na_v_1.8 and is consistently shown to be upregulated in PTHS mouse models. In this perspective, we review prior literature and present novel data that suggests inhibiting Na_v_1.8 in PTHS mouse models is effective at normalizing neuron function, brain circuit activity and behavioral abnormalities and posit this therapeutic approach as a treatment for PTHS.

## Introduction

Pitt Hopkins Syndrome (PTHS) is a rare neurodevelopmental disorder resulting from autosomal dominant mutations on chromosome 18 at the *TCF4* (also known as ITF2, SEF2, E2-2, not <u>T</u>-<u>c</u>ell factor <u>4</u> which is encoded by TCF7L2 gene) locus. Disease-causing mutations are primarily de novo with rare instances of parental mosaicism (1,2) and result in TCF4 haploinsufficiency or dominant negative mechanisms (3–7). PTHS patients display features of ASD and are more generally characterized by intellectual disability, developmental delay, breathing abnormalities, absent or limited speech, motor delay, seizure, constipation, and facial features including wide mouth and a broad nasal base with high bridge (8–11). Exactly how mutations in *TCF4* lead to this disorder remains an open question. However, several studies using PTHS animal models have identified a variety of phenotypes that provide important biological insights into this disorder. These phenotypes are observed across the lifespan, beginning with alterations in cortical development, cell fate specification, neuron development and eventually lead to altered neuronal excitability, synaptic plasticity, and behavioral deficits in adult mice (12–14). Here, we highlight evidence that suggests mutations in *Tcf4* lead to ectopic expression of *Scn10a*/Na_v_1.8 which partially underlies neuronal excitability, network synchronicity and behavioral deficits observed in PTHS mouse models. Moreover, we discuss evidence that inhibition of Na_v_1.8 is effective at acutely rescuing these phenotypes and discuss the potential of Na_v_1.8 as a therapeutic target for the treatment of PTHS.

### Identification of SCN10a/Na_v_1.8 in PTHS

*Scn10a*/Na_v_1.8 was first identified as a downstream dysregulated gene of *Tcf4* in a rat model of PTHS (6). In this model system, shRNA and CRISPR/Cas9 constructs specific to *Tcf4* were delivered by in utero electroporation leading to cellular transgenesis of layer 2/3 pyramidal neurons and knockdown of *Tcf4*. This knockdown resulted in a significant reduction in the intrinsic excitability of transfected neurons. Molecular profiling of transfected neurons via translating affinity purification (iTRAP) led to the identification of two upregulated ion channel genes, *Scn10a* and *Kcnq1*. Rescue experiments with antagonists to these two channels and phenocopy experiments via overexpression of *Scn10a* in wildtype neurons validated the causal role of *Scn10a* and *Kcnq1* in these intrinsic excitability deficits. Further confirmation of the TCF4-dependent excitability deficits were obtained in two different PTHS mouse models (6,15). In the *Tcf4*^*+/tr*^ mouse model, it was shown that SCN10a expression was upregulated, and consistent with the rat model, pharmacological blockade of Na_v_1.8 normalized intrinsic excitability deficits (6). Regulation of *Scn10a* by *Tcf4* appears to be direct, as TCF4 ChIP-seq analysis in rat neuroprogenitor cell cultures indicated that *Tcf4* binds directly to regions of the *Scn10a* genetic locus and therefore is predicted to act as a repressor of *Scn10a* gene expression in the central nervous system (CNS)(6). Together, these initial findings indicated Na_v_1.8 was dysregulated in PTHS rodent models and that its ectopic expression was a key molecular mechanism underlying TCF4-dependent intrinsic excitability deficits. Fortunately, the unique properties of Na_v_1.8 make it a suitable drug target.

### SCN10a/Na_v_1.8 function and pharmacology

SCN10a/Na_v_1.8 is a primarily peripherally expressed, TTX resistant, voltage-gated sodium channel (16), but its expression and function in the central nervous system is reported (6,17,18) and SCN10a variants are associated with epileptic disorders (19). In the peripheral nervous system, Na_v_1.8 is thought to play an important role in nociception (20–23) and in dorsal root ganglion cells (DRGs) Na_v_1.8 is responsible for a substantial proportion of the inward current needed to generate an action potential (24). In addition, Na_v_1.8 also appears to regulate the frequency of action potential firing and spike-frequency adaptation due to its unique kinetic properties (25,26). Na_v_1.8 channels display prominent slow inactivation (16) and DRGs show a pronounced adaptation of action potential firing in response to stimulation (26). Selective inhibitors of Na_v_1.8 have been developed and have shown promise in rodent pain models as well as in early phase human trials. The selective Na_v_1.8 inhibitor A-803467 has shown significant effects on the maximal amplitude and kinetic properties of the TTX-resistant sodium current in rats (17). A-803467, exhibited high affinity and selectivity for blocking human Na_v_1.8 channels and effectively inhibited spontaneous and evoked DRG neuronal action potentials *in vivo* in rats. A-803467 also dose-dependently reduced nociception in neuropathic and inflammatory pain models (21). However, A-803467 in preclinical models has limited oral bioavailability (27). PF-04531083 was developed as a potent and highly selective Na_v_1.8 inhibitor with acceptable oral bioavailability and showed effectiveness in preclinical pain models (28). Moreover, PF-04531083 can pass the blood brain barrier as it was shown to rescue CNS phenotypes in a PTHS mouse model (18). More recently, VX-548 an oral selective Na_v_1.8 inhibitor has shown success in two phase 2 clinical trials for acute pain in patients who had recently undergone abdominoplasty or bunionectomy (29,30), however the ability of VX-548 to penetrate the blood brain barrier is not known.

### Normalization of breathing and behavioral abnormalities

A common symptom observed in PTHS patients is disordered breathing characterized by hyperventilation and intermittent apnea or breath holding (31,32). These breathing abnormalities severely impact the patient’s quality of life and often contribute to aspiration-induced pneumonia, which is the leading cause of death in PTHS (33,34). Remarkably, similar breathing abnormalities were observed in a PTHS mouse model (18). *Tcf4*^*+/tr*^ mice display frequent episodes of hyperventilation, reduced sigh activity, increased post-sigh apnea, and fail to increase inspiratory and expiratory output in response to CO_2_. Cleary and colleagues deduced that these breathing abnormalities may result from abnormal function of the retrotrapezoid nucleus (RTN) because similar breathing abnormalities are found in Rett Syndrome and are known to involve chemoreception. In addition, acetazolamide, a carbonic anhydrase inhibitor, used to induce metabolic acidosis and hyperventilation, improved breathing in PTHS patients (35–40). They showed that *TCF4* mutation resulted in selective loss of parafacial Phox2b+ neurons, altered connectivity between Phox2b+ neurons and the pre-BotC complex, and suppressed excitability of chemosensitive RTN neurons. All these phenotypes were consistent with previously observed phenotypes in various brain regions of PTHS mouse models (6,15,41,42). They went on to show that *Scn10a* expression is not normally detected in the RTN of WT mice, however *Scn10a* expression was observed in *Tcf4*^*+/tr*^ mice, and pharmacological block of Na_v_1.8 with IP injection of PF-04531083 was effective at rescuing breathing in these animals. Moreover, they showed that acute Na_v_1.8 block was also effective at rescuing hyperlocomotion and anxiety in the *Tcf4*^*+/tr*^ mice. Importantly, they demonstrated that rescue by PF-04531083 was specific to inhibition of Na_v_1.8 in the CNS, because IP injection of PF-06305591, which does not penetrate the blood brain barrier, was ineffective at normalizing behavior.

Together, Cleary and colleagues provided direct *in vivo* evidence showing that central inhibition of Na_v_1.8 was effective at normalizing breathing and behavioral abnormalities in a PTHS mouse model, further supporting the idea of Na_v_1.8 as a therapeutic target. In another set of studies, Ekins and colleagues performed a high throughput screen to identify FDA approved drugs for inhibition on recombinant Na_v_1.8 expressed in HEK cells (43). Their screen identified a number of dihydropyridine calcium channel antagonists that were effective at blocking Na_v_1.8 channels, with nicardipine being the most potent with a sub micromolar IC50 (0.6uM). They went on to show that administration of nicardipine improved several behavioral deficits in a PTHS mouse model, including social recognition, nesting, self-grooming, fear conditioning, and hyperlocomotion (43). However, the exact mechanism of rescue by nicardipine is not entirely clear, as it is likely inhibiting both sodium and calcium channels. Overall, these studies provide evidence that inhibition of Na_v_1.8 is effective at rescuing breathing and behavioral abnormalities in PTHS mouse models and therefore support therapeutic targeting of Na_v_1.8.

### Normalization of auditory evoked potentials

Event-related potentials (ERPs) are stereotyped patterns of voltage fluctuation measured in response to sensory stimuli, which consist of temporal components that reflect physiological response. Levels of spectral power and phase coherence during ERP components are thought to reflect strength and connectivity in cortical circuits that mediate sensory information processing (44). Following our previously published methods (45), we recorded auditory ERPs in wild-type (WT) and *Tcf4*^*+/t*r^ mice at baseline (vehicle) and after acute administration of the Na_v_1.8 antagonist PF-04531083 (10mg/kg, i.p.). We used component and time-frequency analysis of the ERP to identify changes in patterns of synchronized oscillatory activity during the ERP in this PTHS mouse model at baseline and following Na_v_1.8 antagonism. Component analysis of the ERP showed that there is a significant effect of genotype in reducing the N40 amplitude peak (Figure 1B-D). In addition, the event-related spectral perturbation (ERSP) power showed changes relative to tone onset in *Tcf4*^+/tr^ mice and alterations in phase locking as measured by intertrial phase coherence (ITC, Figure 2). Specifically, we observed no difference in ERSP at baseline between genotypes (Figure 1A and data not shown) but observed significant delay in the latency of low (theta) frequency activity (Figure 1E), and increased level of coherence in the high (gamma) frequency ITC at baseline (Figure 1D). The delayed oscillatory activity and increased gamma synchrony in response to auditory stimuli suggests impairments in the neural correlates of sensory information processing in this PTHS mouse model. Given prior evidence that Na_v_1.8 is upregulated in this mouse model and that Na_v_1.8 antagonists were effective at normalizing both intrinsic excitability and behavior, we quantified the acute effect of PF-04531083 on ERPs. PF-04531083 had no effect on N40 peak amplitude in WT or *Tcf4*^+/tr^ mice (Figure 1B-D). However, PF-04531083 did significantly reduce gamma ERSP in *Tcf4*^+/tr^ mice, but not in WT mice (Figure 2A, C). In addition, acute Na_v_1.8 blockade also normalized the latency of theta ITC and gamma ITC (Figure 2B, D, E). These results suggest acute Na_v_1.8 antagonism is effective at normalizing abnormal synchronous activity in the PTHS mouse model and provides further support that Na_v_1.8 may have utility for treating symptoms in PTHS. Moreover, these data represent a potential electrophysiological biomarker that could be utilized for screening Na_v_1.8 antagonists for therapeutic efficacy. Moreover, patterns of oscillatory activity are well-conserved across species, and if similarly altered ERP responses were detected by scalp EEG recordings in PTHS patients, the translational value of this biomarker would be invaluable.

**Figure 1.**
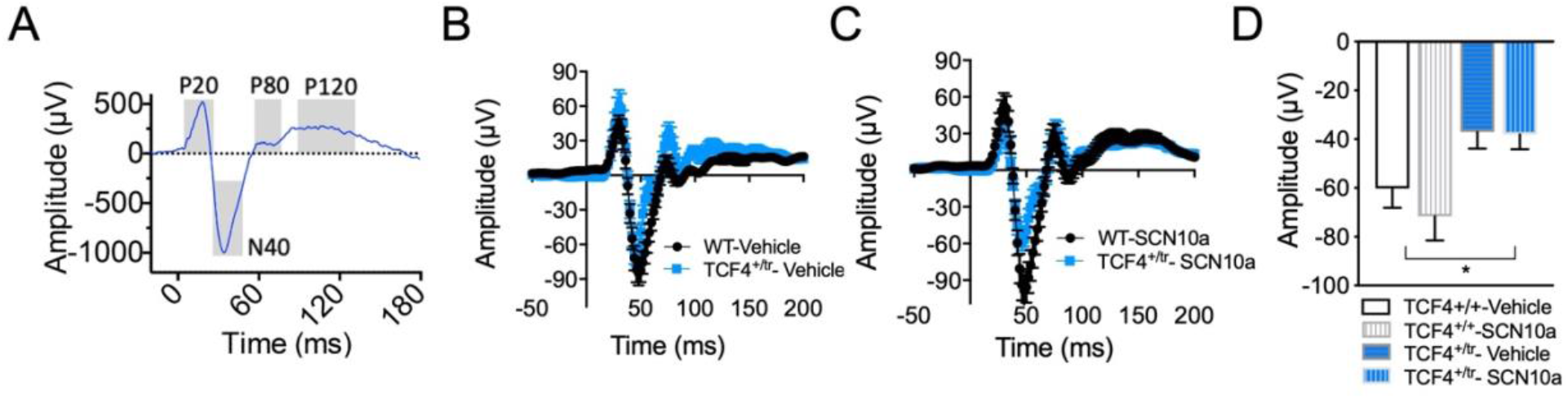
Sensory information processing deficits are normalized by Na_v_1.8 inhibition. **(A)** Example event-related potential (ERP) grand averages from individual temporal components (P20, N40, P80 and P120) where time 0=auditory stimulus (S1) onset. **(B)** Grand average ERPs in *Tcf4*^*+/tr*^ (n=10) compared to WT (n=12) animals at baseline and (C) following PF-04531083 administration. **(D)** Summary component analysis showing significantly reduced amplitudes in N40 peaks in *Tcf4*^*+/tr*^ mice compared to WT animals, which are not altered by PF-04531083 administration (2-way RM ANOVA, * p=0.0163, main effect of genotype; ns p=0.1317, main effect of treatment).

**Figure 2.**
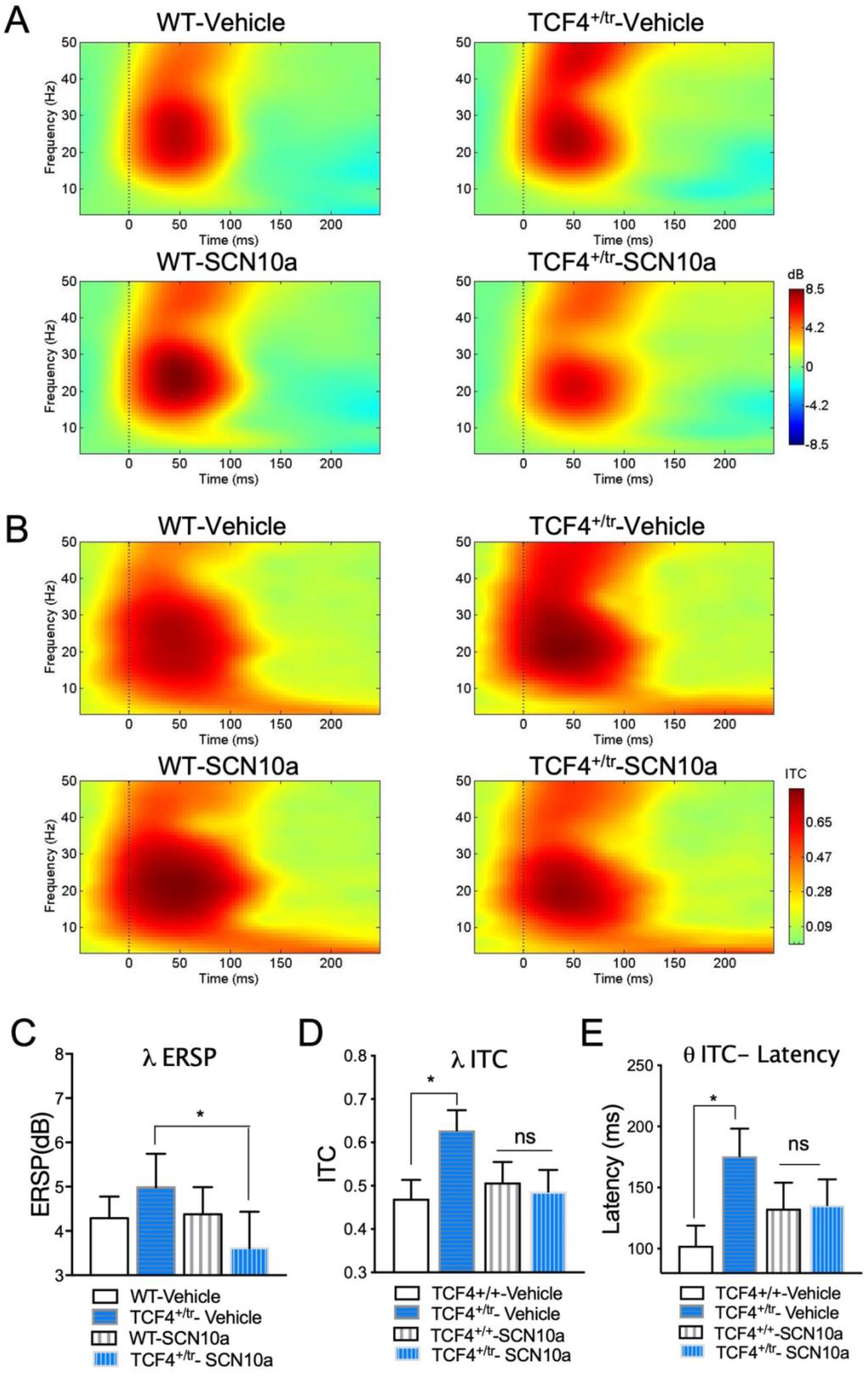
**(A)** Heat maps of event-related spectral perturbation (ERSP) in WT (left, n=12) and *Tcf4*^*+/tr*^ (right, n=10) animals depicting ERP-related changes due to genotype (vehicle) and rescue with PF-04531083 (SCN10a). **(B)** Heat maps of intertrial coherence (ITC) in WT (left, n=12) and *Tcf4*^*+/tr*^ (right, n=10) animals depicting ERP-related changes due to genotype (vehicle) and rescue with PF-04531083 (SCN10a). **(C)** Reduction of gamma ERSP following SCN10a antagonism in *Tcf4*^*+/tr*^, but not in WT animals (2 way RM ANOVA, p=0.0453 interaction of genotype X treatment; Bonferroni post hoc, *p=0.0269 vehicle-treated *Tcf4*^*+/tr*^ versus PF-04531083-treated *Tcf4*^*+/tr*^; ns p>0.9999 vehicle-treated WT versus PF-04531083-treated WT). **(D)** High frequency disturbances in *Tcf4*^*+/tr*^ mice are corrected by SCN10a antagonist. There is significantly higher gamma ITC in vehicle-treated *Tcf4*^*+/tr*^ compared to WT vehicle-treated mice in the first 75 ms post-tone. Following SCN10a treatment, there is no effect of genotype (2 way RM ANOVA, p=0.0027 interaction of genotype X treatment; Bonferroni post hoc, *p=0.0446 vehicle-treated WT versus *Tcf4*^*+/tr*^; ns p>0.9999 PF-04531083-treated WT versus *Tcf4*^*+/tr*^). **(E)** Low frequency disturbances in *Tcf4*^*+/tr*^ mice are corrected by PF-04531083. Latency to peak theta (3-8 Hz) ITC is significantly increased in vehicle-treated *Tcf4*^*+/tr*^ compared to vehicle-treated WT mice. No significant effect of genotype is detected following treatment with the SCN10a antagonist (2 way RM ANOVA, p=0.0123 interaction of genotype X treatment; Bonferroni post hoc, *p=0.0303 vehicle-treated WT versus *Tcf4*^*+/tr*^; ns p>0.9999 PF-04531083-treated WT versus *Tcf4*^*+/tr*^).

### SCN10a/Na_v_1.8 and myelination

Another potential therapeutic benefit of Na_v_1.8 antagonists in PTHS could be through its relation to demyelinating disorders. Transcriptional profiling of several different PTHS mouse models showed that differentially expressed genes were enriched in neurons and oligodendrocytes (OLs), and analysis of OLs and myelination in the *Tcf4*^*+/tr*^ mouse showed a significant reduction in OL density, myelination and function (41). These results suggest that re-myelination could be a potential therapeutic avenue for PTHS but may also provide another link to therapeutic targeting of *Scn10a*/Na_v_1.8. Several groups have shown that a variety of diseases associated with demyelination result in maladaptive ectopic expression of *Scn10a*/Na_v_1.8. For instance, hereditary demyelinating neuropathy leads to an upregulation of *Scn10a*/Na_v_1.8 and abnormal axonal excitability (46), and ectopic *Scn10a*/Na_v_1.8 is observed in the cerebellum of the experimental autoimmune encephalomyelitis (EAE) mouse model of multiple sclerosis (MS) and in MS patients (47). These results have led to the notion that Na_v_1.8 antagonists may be a beneficial treatment for demyelinating diseases and neuropathies (23). It was subsequently shown that an administration of an orally bioavailable Na_v_1.8 antagonist (PF-01247324) improved cerebellar-dependent motor coordination in a transgenic mouse model overexpressing *Scn10a* as well as the EAE mouse model of MS (48,49). The link between demyelination and *Scn10a* expression is intriguing, and a similar maladaptive mechanism could be at play in PTHS in response to the TCF4-dependent reduction in myelination. Overall, these results suggest inhibition of Na_v_1.8 in PTHS patients may provide a dual benefit by normalizing neuronal excitability and improving myelin related deficits.

## Conclusion

Currently there are no approved medications for the core symptoms of ASD or even subsets of ASD like PTHS. Here, we discuss the results of a variety of rodent studies on PTHS that all converge on Na_v_1.8 as being a plausible therapeutic target. Rodent models of PTHS have routinely shown that disruption of *Tcf4* function leads to upregulation of *Scn10a*/Na_v_1.8 and pharmacological blockade of Na_v_1.8 is effective at normalizing both physiological and behavioral phenotypes. Potent and selective Na_v_1.8 antagonists are developed and their safety in humans is demonstrated in clinical trials (50,51). Given all these factors, we recommend testing antagonists of Na_v_1.8 as a therapeutic approach for PTHS.

## Acknowledgements

We are grateful for the vision and generosity of the Lieber and Maltz families, who made this work possible. This work was supported by the Lieber Institute for Brain Development, the Pitt–Hopkins Research Foundation Awards (to B.J.M.), National Institute of Mental Health (NIMH) grant R56MH104593 (to B.J.M.), NIMH grant R01MH110487 (to B.J.M.). We thank Julia Hill for performing and analyzing experiments.

## Author contributions

K.M. and B.J.M designed the experiments. K.M analyzed the results. D.D. S.R.S. and B.J.M. wrote the manuscript. K.M. and B.J.M reviewed and edited the manuscript.

